# Using breathing systems in anaesthesia for up to 7 days instead of 24 hours: a comparative microbial safety study

**DOI:** 10.1101/2024.07.11.603054

**Authors:** Cynthia P. Haanappel, Elisabeth A. Rieff, Lucija Pavkovic, Merel N. van Holst-Raaphorst, Woutrinus de Groot, Caroline D. van der Marel, Anne F. Voor in ’t holt, Juliëtte A. Severin

## Abstract

**Summary:** The replacement frequency of mechanical ventilator’s breathing systems used in operating rooms (ORs) currently varies between hospitals. In light of evidence-based decision-making and sustainability efforts, we aim to determine whether 7-day use of breathing systems instead of 24 hours is microbial safe. In this prospective single-centre explorative study, 30mm UniflowTM breathing systems used in eight ORs were included. In four ORs, breathing systems were replaced daily following standard practice. In the remaining four ORs, they were intended for a 7-day use. Breathing systems were sampled daily on three locations of the exterior surface and cultured for the presence of microorganisms. A total of 128 breathing systems were included, 99 from an OR with daily replacement and 29 from an OR with weekly replacement. A total of 604 samples were cultured, of which the majority, 549 (90.9%) cultures were negative. From the 55 (9.1%) positive cultures, the majority (n=49, 70%) were coagulase-negative staphylococci. None of the identified microorganisms were found in consecutive cultures. Cultures from day 2 to 7 did not show a statistically significant increased positivity rate compared to cultures from day 1, respectively 22.9% vs. 24.1%. The weekly replacement regimen, furthermore, decreased the number of breathing systems used with 71%.

Our data indicates that use of breathing systems up to seven days remains microbial safe. Additionally, only a minimal number of pathogenic microorganisms were detected, and these were not persistent on the breathing systems. Transitioning from 24-hours to intended 7-day use could significantly reduce costs and CO_2_ emissions.

## Introduction

Anaesthetic breathing systems are crucial for the delivery of mechanical ventilation of patients undergoing surgical procedures conducted in the operating room (OR). These systems may be used for more than one patient, and even up to seven days according to manufacturer’s instructions. This practice is based on internationally approved and certified breathing circuit filters, which protect the circuit from contamination and thus prevent the risk of pathogen transmission via the tube system’s lumen [1, 2]. However, this approach does not consider the potential contamination of the exterior surfaces of the breathing systems. In many hospitals worldwide, breathing systems are replaced after every patient or on a daily basis. Frequent replacement is based on the general concept that inanimate surfaces in healthcare settings may become contaminated with pathogens, presenting a potential risk for nosocomial infections, including surgical site infections (SSI) [3, 4]. Cleaning (and disinfection) can overcome this potential risk, however, this is not possible for the breathing systems due to the ribbed tube surface. This is also the reason why the systems are always replaced after surgery in a patient who is known to carry a multidrug-resistant microorganism (MDRO) or another microorganism for which additional cleaning regimens are required, or when the system is visually contaminated.

In light of evidence-based decision-making and reducing hospital waste [5], we questioned if adapting the daily replacement of breathing systems protocol to once a week could reduce waste, without affecting patient safety. In several hospitals, breathing systems are already used for seven days, and it is even questioned if they can be used for up to 4 weeks [6, 7]. Besides a reduction in CO_2_ emission, it reduces workload for healthcare workers and has economic advantages. Therefore, this study aims to determine whether using breathing systems in anaesthesia for up to 7 days instead of 24 hours, leads to an increase of contamination of the exterior surface with possible pathogenic microorganisms over time.

## Methods

This was a prospective single-centre explorative study, performed at the Erasmus MC University Medical Centre in Rotterdam, The Netherlands. The Erasmus MC is a tertiary care centre with 30,000 yearly patient admission, 39 ORs and 25,000 yearly surgical procedures. The anaesthetic breathing systems used in our hospital are 30mm Uniflow™ breathing systems.

For this study, we included breathing systems from eight different ORs, which were mainly used for procedures by the departments of general surgery, neurosurgery, and ear nose and throat (ENT). In four ORs, breathing systems were replaced daily, following standard practice. In the remaining four ORs, they were intended for a 7-day use, with a standard replacement procedure every Monday. Breathing systems in the weekly replacement schedule were prematurely replaced if there was visual contamination, usage of the breathing systems with a patient identified with a MDRO or another microorganism for which isolation and contact precautions were indicated, or if there was wear-and-tear of the system.

Cultures to determine contamination were taken from three different sample locations per culture moment, together defined as a “culture set” (Figure 1). Samples were taken from the location closest to the patient near the attachment to the endotracheal tube, from the mid part of the system and closest to the mechanical ventilation machine. Sampling occurred every day at 4 PM, by an infection prevention and control practitioner or healthcare worker from anaesthetics.

**Figure 1.**
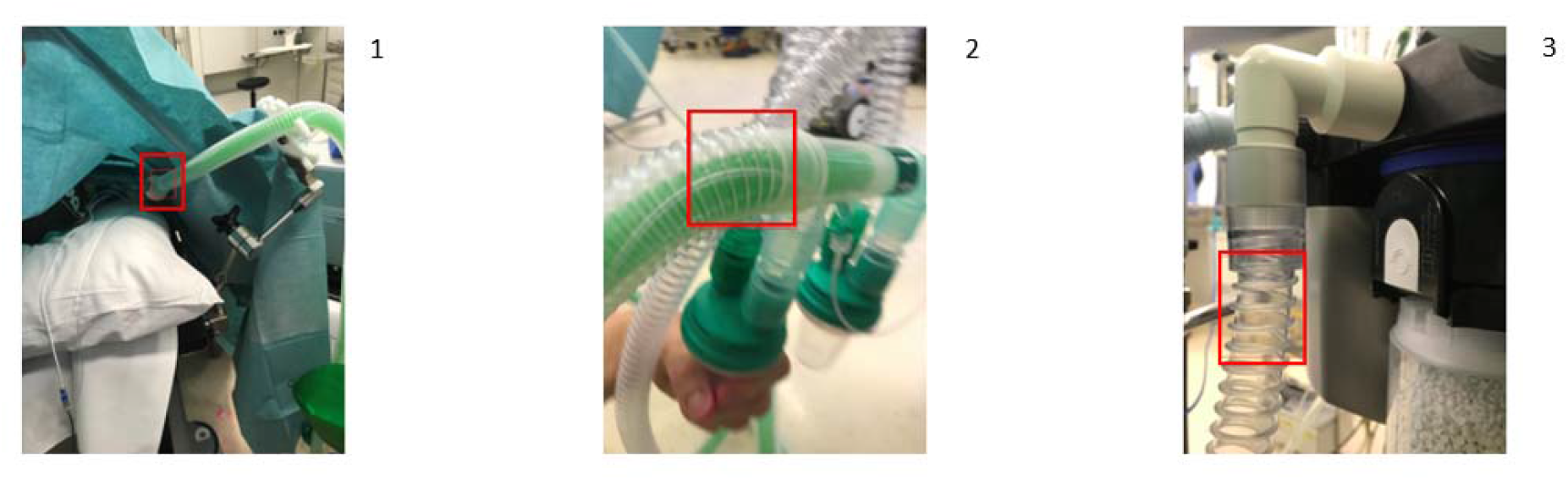
Sample locations of the breathing systems. ^1^Sample location closest to the patient, near the attachment to the endotracheal tube. ^2^Sample location in the mid part of the system. ^3^Sample location closest to the mechanical ventilation machine.

Samples were taken with BD BBL™ CultureSwab™ Plus cotton swabs (Copan Italia, Brescia, Italy) and stored in the accompanying 1 mL Amies gel transport medium. Before sampling, cotton swabs were pre-wetted with phosphate buffered saline (PBS). Swabs were plated directly on blood agar and MacConkey agar plates (bioMérieux, Marcy l’Etoile, France), which were then incubated for 48 hours at 35°C. Bacterial colonies were identified to species level using Matrix-Assisted Laser Desorption/Ionization Time-Of-Flight mass spectrometry (MALDI-TOF [Bruker Daltonics, Bremen, Germany]). Antimicrobial susceptibility was determined for *Staphylococcus aureus, Enterococcus* spp., *Pseudomonas* spp., *Acinetobacter* spp., *Enterobacterales*, and *Stenotrophomonas* spp., using the VITEK®2 system (bioMérieux, Marcy l’Etoile, France). If there was growth in the culture of at least one of the three sample locations, a culture set was considered positive.

Descriptive and explorative analyses were performed, and missing data were described and excluded from further analysis. Categorical variables were displayed in absolute number and percentages. Chi-square tests to test for statistical significance were performed using R version 4.2.1, whereby P<0.05 was considered to be statistically significant.

## Results

A total of 142 breathing systems were eligible for inclusion, of which 128 (90.1%) breathing systems with 206 culture sets were included. Reasons for which 14 breathing systems or culture sets could not be included were among others, that breathing systems were either replaced before the sampling moment could take place, or because the sample location was not accessible due to ongoing surgery during the sampling moment. Of the 128 included breathing systems, 99 (77.3%) were present in an OR with a daily replacement regimen and 29 (22.7%) breathing systems in an OR with a weekly replacement regimen. In total, 604 samples were obtained, 14 samples of 13 culture sets were missing. The majority of cultures 549 (90.9%) were negative. From the 55 positive cultures, 70 different microorganisms were identified, of which the majority (n=49, 70%) were coagulase negative staphylococci (CoNS). All identified species are reported in Supplementary Table A1. The most noteworthy found microorganisms which are potentially pathogenic, were *Staphylococcus aureus, Enterococcus faecium*, and *Acinetobacter* species. None of the found microorganisms were multidrug resistant [8]. Furthermore, all microorganisms were only found once on breathing systems, meaning no species were found in consecutive cultures from breathing systems used for longer than 24 hours. Of the different sample sites, location 1 had the most positive samples with 35 of 195 (17.9%) samples. Sample location 2 and 3 had 15 of 205 (7.3%) and 5 of 205 (2.5%) positive cultures, respectively.

When comparing the overall positivity rate of culture sets from breathing systems in daily replacement regimen OR (27 of 99; 27.3%) to those used in OR with weekly replacement regimen (21 of 107; 19.6%), no statistical significant difference was found (P = 0.195). From all day 1 culture sets, 29 of 127 (22.9%) sets were positive and from all cultures from day 2 to 7, 19 of 79 (24.1%) were positive, this difference was also not statistically significant (P = 0.841). Additionally, no increasing positivity rate per day could be observed from culture sets from breathing systems with weekly replacement (Figure 2).

**Figure 2.**
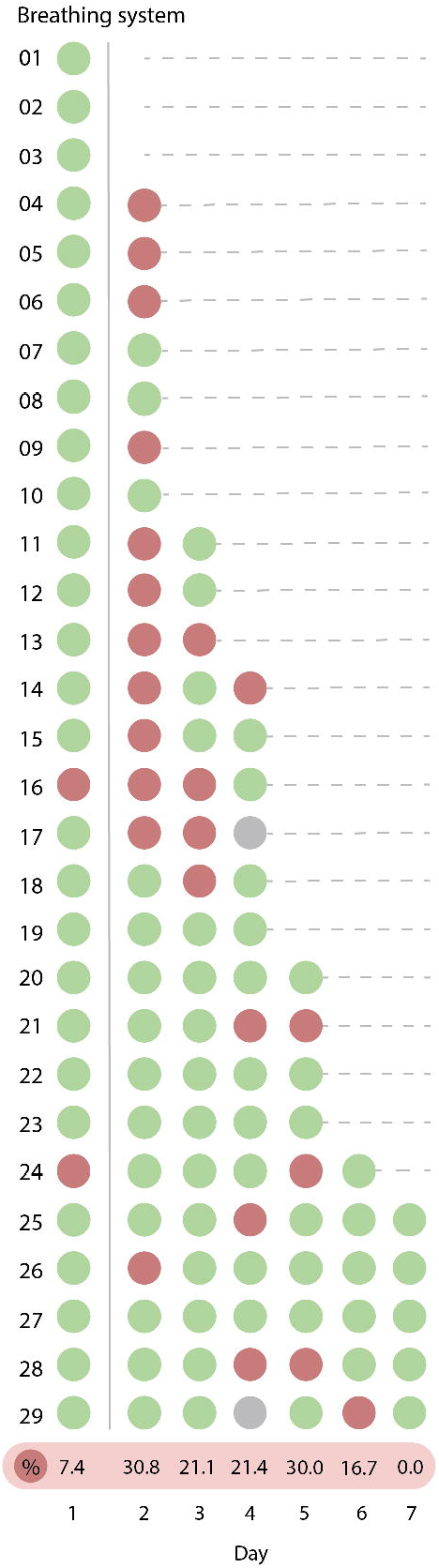
Culture results from breathing systems used in operating rooms with weekly replacement. Green: Negative culture, Red: positive culture, Grey: culture result not available, %: describes the positivity rate of cultures per day of use.

As displayed in Figure 2, the majority of breathing systems used in ORs with a weekly replacement regimen were not used for seven consecutive days. Only five out of 29 breathing systems (17.2%) reached seven days of use. Reasons for premature replacement of breathing systems were; replacement after use with a patient with an isolation label due to MDRO carriage or another microorganism for which isolation measures were indicated (n=6), the breathing system was visibly contaminated (n=5), wear-and-tear of the breathing system (n=1), a different breathing system used (n=1), or unknown (n=1). The remainder of breathing systems were replaced on every Monday on the weekly replacement moment.

Adjusting the breathing system replacement frequency from daily to weekly in the OR can lead to a substantial decrease in waste production and with that a decrease in CO_2_ emission and eventual costs. During our study period, 99 breathing system were used on the ORs with a daily replacement frequency, while the ORs with a weekly replacement frequency used 29 breathing systems. This results in a consumption reduction of 71%. Based on the consumption of breathing systems in 2022, a year when the OR production was almost back to pre-COVID-19 level, an estimation was made from the potential cost- and CO_2_-savings when the current daily replacement policy would change to weekly. This lower replacement frequency could result in a yearly reduction of around 3000 Kg plastic and 12 000 Kg CO_2_ emissions (Table I).

**Table I.** Estimated CO_2_- and cost reduction based on yearly consumption of breathing systems.

|  | Replacement frequency |  | 71% reduction per year | Savings (71% reduction – weekly replacement) |
| --- | --- | --- | --- | --- |
|  | Daily* | Weekly |  |  |
| Breathing systems used (N) | 5520 | 1104 | 1601 | 3919-4416 |
| Costs (€)(€20 per breathing system) | 110400 | 22080 | 32020 | 78380-88320 |
| Plastic in Kg (0.76Kg/breathing system) | 4195 | 839 | 1217 | 2978-3356 |
| Environmental impact in Kg CO<sub>2</sub>** | 15522 | 3104 | 4501 | 11021-12417 |

\* on working days = 5 breathing systems per week
\*\* 3.7Kg CO<sub>2</sub>eq/kg plastic [9], whereby 1 kg CO<sub>2</sub>-equivalent is equal to the greenhouse effect of 1 kg CO<sub>2</sub>.

## Discussion

The findings of this study do not indicate an increase in the exterior surface contamination of breathing systems with potential pathogenic microorganisms, when used up until seven days without intermediate cleaning procedures. Even though the number of breathing systems used up until seven days in our study was limited, our results do not provide any indication that use of these systems up to seven days creates a microbial unsafe situation. This is mainly due to the predominant presence of commensal skin flora (CoNS) among the microorganisms identified in the samples. Additionally, only a minimal number of pathogenic microorganisms were detected, and these were not persistent on the breathing systems.

A possible explanation for the limited contamination of the breathing systems might be the already installed infection prevention and control (IPC) measures and care bundle at the OR, including the presence of a plenum (OR air supply system), uniforms and other personal protective equipment, and behaviour of healthcare workers, such as the hand hygiene compliance and number of door openings during surgical procedure [10, 11]. Because of the wide scale of IPC measures in place, which are specifically tailored for the OR, we cannot extrapolate these results to locations or situations outside of the OR complex.

A similar study, however, from Dubler, et al. investigated VentStar® breathing systems and found contamination rates of 6.9% on day 1 versus 16.8% on day 7 on the outer surface of breathing systems whereby the majority of positive culture were also CoNS [12]. One other study reported findings within the same range, with an outer surface contamination of 12.0% on day 1 and 19.4% on day 7, for breathing systems of Tyco® 300/13324 [13]. It is possible that the level of contamination is influenced by the breathing system’s material which makes the results dependent on the brand of breathing systems used. Therefore, no conclusions can be made about similar breathing systems of other brands and their use up until seven days.

Moreover, within this study, we observed that breathing systems remain in use for seven days only in limited numbers. This is because of other reasons for replacement like patients for which isolation and contact precautions are indicated, visible contamination or the wear-and-tear of the breathing systems. Considering that in practice not all breathing systems will be used for seven days, transitioning from 24-hours to intended 7-day use could significantly reduce costs and CO_2_ emissions from waste. The reduction in the amount of plastic and the associated CO_2_ emissions is considerable, although it is based on a rough estimate. For a more accurate estimation, a Life Cycle Analysis (LCA) should be conducted for this specific product and practicalities around pre-mature replacement of breathing systems should be taken into account. Nevertheless, reducing waste by prolonging the replacement frequency will contribute to making healthcare more sustainable on a large scale without increasing the risk of infection for individual patients [5]. To provide more evidence-based sustainable IPC programmes, more research is needed to define microbial safety further. In this instance, a randomized controlled trial is not feasible, but other research methodologies might provide further input for better evidence-based decision-making. Furthermore, it could be worthwhile to investigate the use of breathing systems for longer than seven days.

## Conclusion

Our data indicate that prolonged use of breathing systems up to seven days can remain microbial safe. Whether even longer periods of use can be applied, will depend on the manufacturer’s instructions and limitations in use, but could be subject of further research.

## Funding

None.

## Ethical approval

This study was approved by the medical ethical committee of the Erasmus MC (MEC-2023-0586) and was not subject to the Medical Research Involving Human Subjects Act.

## Conflict of interest statement

None declared.

## Supplementary Materials

**Supplementary Table A1**. Identified microorganism species on breathing systems.

## Notes

### Competing Interest Statement

The authors have declared no competing interest.

