## Supplementary Table A1 for "Using breathing systems in anaesthesia for up to 7 days instead of 24 hours: a comparative microbial safety study"

**Supplementary Materials**

**Supplementary Table A1.** Identified microorganism species on breathing systems.

| **Microorganisms** | **N** | **%** |
| --- | --- | --- |
| *Staphylococcus hominis* (CoNS) | 13 | 18.6% |
| *Staphylococcus epidermidis* (CoNS) | 13 | 18.6% |
| *Staphylococcus capitis* (CoNS) | 10 | 14.3% |
| *Staphylococcus aureus* | 4 | 5.7% |
| *Staphylococcus warneri* (CoNS) | 3 | 4.3% |
| *Staphylococcus haemolyticus* (CoNS) | 3 | 4.3% |
| *Bacillus cereus* group | 2 | 2.9% |
| *Streptococcus mitis* group | 2 | 2.9% |
| *Micrococcus luteus* | 2 | 2.9% |
| *Staphylococcus pasteuri* (CoNS) | 2 | 2.9% |
| *Staphylococcus pettenkoferi* (CoNS) | 2 | 2.9% |
| *Enterococcus faecium* | 1 | 1.4% |
| *Corynebacterium coyleae* | 1 | 1.4% |
| *Staphylococcus epidermidis* (CoNS) | 1 | 1.4% |
| Gram-positive cocci (clusters) | 1 | 1.4% |
| *Rothia mucilaginosa* | 1 | 1.4% |
| *Neisseria* species | 1 | 1.4% |
| *Kytococcus* species | 1 | 1.4% |
| *Rothia koreensis* | 1 | 1.4% |
| *Acinetobacter* species | 1 | 1.4% |
| *Streptococcus infantis* (*S. mitis* groep) | 1 | 1.4% |
| *Brevibacterium* species | 1 | 1.4% |
| Viridans streptococci | 1 | 1.4% |
| *Staphylococcus caprae* (CoNS) | 1 | 1.4% |
| *Staphylococcus cohnii* (CoNS) | 1 | 1.4% |
